# Environmental RNA as a non-invasive tool for assessing toxic effects in fish: a proof-of-concept study using Japanese medaka exposed to pyrene

**DOI:** 10.1101/2023.04.25.538202

**Authors:** Kyoshiro Hiki, Takahiro Yamagishi, Hiroshi Yamamoto

**Author notes:** **Author contributions** Kyoshiro Hiki: Conceptualization, Methodology, Investigation, Formal analysis, Writing – original draft. Takahiro Yamagishi: Conceptualization, Methodology, Writing – review & editing. Hiroshi Yamamoto: Supervision, Resources, Writing – review & editing.

## Abstract

Although environmental RNA (eRNA) is emerging as a non-invasive tool to assess the health status of aquatic macro-organisms, the potential of eRNA remains largely untested. In this study, we investigated the ability of eRNA to detect changes in gene expression in Japanese medaka fish (*Oryzias latipes*) in response to sub-lethal pyrene exposure, as a model toxic chemical. We performed standardized acute toxicity tests and collected eRNA from tank water and RNA from fish tissue after 96 h of exposure. Our results showed that over 1000 genes were detected in eRNA and the sequenced read counts of these genes correlated with those in fish tissue (*r* = 0.50). Moreover, eRNA detected 86 differentially expressed genes in response to pyrene, some of which were shared by fish RNA, including suppression of collagen fiber genes. These results suggest that eRNA has the potential to detect changes in gene expression in fish in response to environmental stressors without the need for sacrificing or causing pain to fish. However, we also found that the majority of sequenced reads of eRNA (> 99%) were not mapped to the reference medaka genome and they originated from bacteria and fungi, resulting in low sequencing-depth. In addition, eRNA, in particular nuclear genes, was highly degraded with a median transcript integrity number (TIN) of < 20. These limitations highlight the need for future studies to improve the analytical methods of eRNA application.

**TOC Art:** 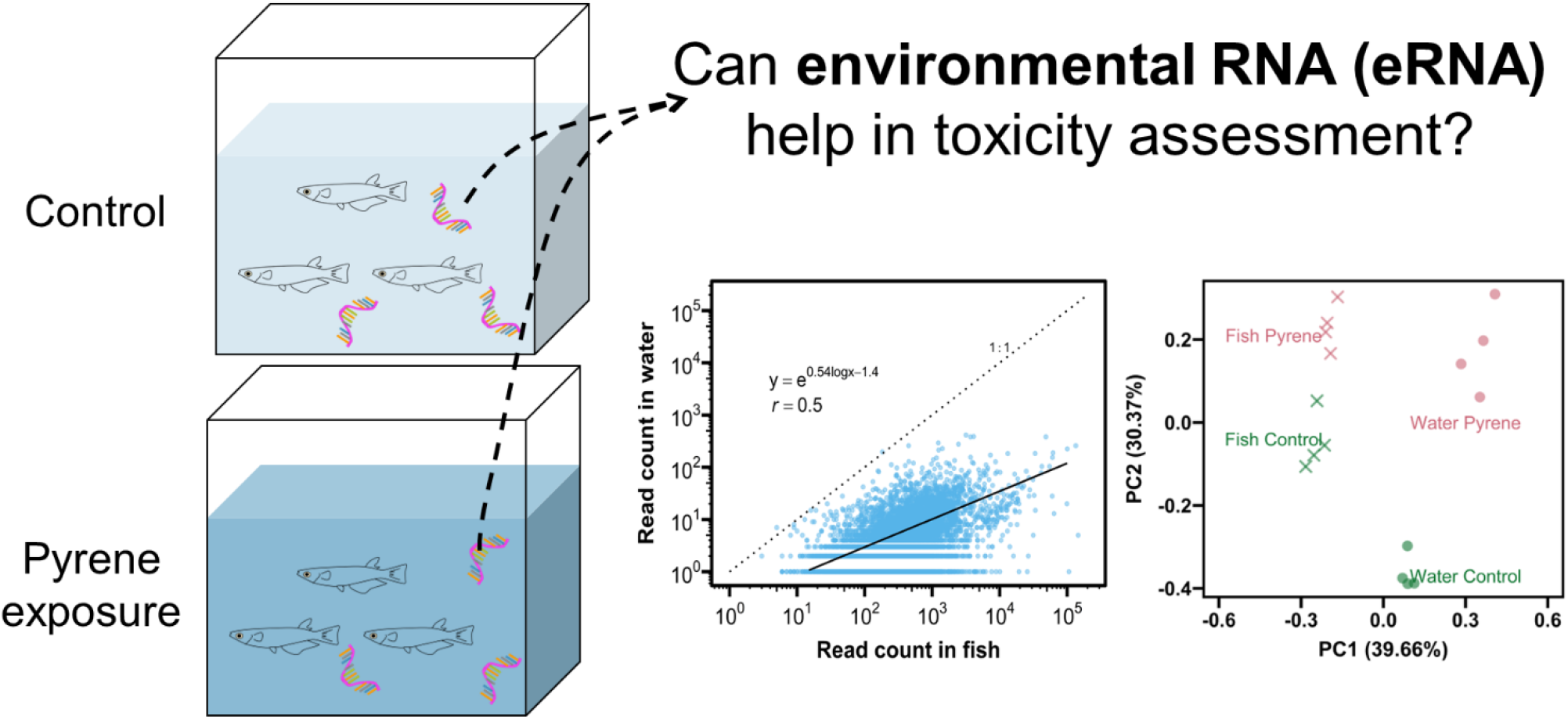

## 1. INTRODUCTION

Toxicity testing using aquatic animals have been widely conducted for chemical risk assessments. While collecting and analyzing tissues or blood samples from animals exposed to test chemicals can provide valuable insights into the molecular mechanisms of toxicity, this process can also cause harm or death to the animals. In light of increasing concern for animal welfare and ethical issues, some countries have introduced regulations that encourage or require the reduction of animal use, replacement with alternative methods, and refinement of testing procedures to avoid painful or distressing situations (Katsiadaki et al. 2021; Paparella et al. 2021; Yamagishi et al. 2022).

To this end, several non-invasive methods have been proposed to study toxic effects in experimental animals, using materials such as feces (Hano et al. 2021; Liu et al. 2021), fish skin mucus (Meucci and Arukwe 2005; Ekman et al. 2015; Breacker et al. 2017; Oliveira et al. 2018), and fish scales (Alves et al. 2016). These materials contain messenger RNA (mRNA) (Liu et al. 2021), proteins (Meucci and Arukwe 2005; Song et al. 2017), metabolites (Ekman et al. 2015; Hano et al. 2021), or intestinal microbiota (Hano et al. 2021), which can provide information about the physiological status of the experimental animals and reflect their toxic effects. Additionally, traditional behavioral assessments is useful for interpretation of toxicity mode of action (McKim et al. 1987). However, these non-invasive methods are only applicable to a limited range of species. For example, it can be difficult to collect enough feces from small animals (e.g., water fleas) and to collect mucus from species other than fish and amphibians.

Environmental RNA (eRNA) is RNA released by organisms to their surrounding environment such as water, soil, and sediment (Cristescu 2019). Unlike DNA, RNA is only produced for genes that are actively being transcribed, making it a useful tool for understanding the “real time” status of the organisms. Recent studies reported that the degradation of eRNA for some genes was slower than previously expected (Wood et al. 2020; Marshall et al. 2021; Jo et al. 2022; Kagzi et al. 2022). eRNA has been detected for a wide range of aquatic species, including fish (Tsuri et al. 2020; Miyata et al. 2021; Jo et al. 2022), water fleas (Hechler et al. 2022; Kagzi et al. 2022), polychaetas (von Ammon et al. 2019; Wood et al. 2020), mussels (Marshall et al. 2021), ascidians (Wood et al. 2020), and algae (Miyata et al. 2022). Also, eRNA has been detected for both mitochondrial and nuclear genes (Tsuri et al. 2020; Marshall et al. 2021; Jo et al. 2022) and for several microRNAs (Ikert et al. 2021). In addition, eRNA may originate from various tissues including gills, skin, and mucus (Tsuri et al. 2020; Ikert et al. 2021). These findings suggest the potential of eRNA to evaluate the toxic effects in a wide range of organisms exposed to a chemical; however, this potential has yet to be tested.

To explore the potential of eRNA as a non-invasive method for assessing the physiological status of the experimental organisms, we performed toxicity tests using Japanese medaka *Oryzias latipes* as a model fish. Japanese medaka is widely used in ecotoxicity tests for chemical risk assessments (Watanabe et al. 2017; Yamagishi et al. 2022) and has its whole genome published (Kasahara et al. 2007). In addition, as the sex-determining gene DMY has been identified (Matsuda et al. 2002), this species is well-suited for assessing the effects of endocrine-disrupting chemicals (OECD 2015; Watanabe et al. 2017). In this study, the fish were exposed to pyrene, a model toxicant, for 96 h, after which eRNA in the surrounding water was collected, extracted, and comprehensively sequenced. The eRNA profiles were then compared to transcriptome profiles of whole fish tissue. Comprehensive sequencing of eRNA has previously been reported only for the water flea *Daphnia pulex* (Hechler et al. 2022) and rainbow trout *Oncorhynchus mykiss* (Ikert et al. 2021), making this study potentially valuable in understanding eRNA.

## 2. MATERIALS AND METHODS

### 2.1 .Test Organisms and sample preparation

Japanese medaka *O. latipes* was obtained from a brood stock maintained for more than 15 years at the National Institute for Environmental Studies, Japan (NIES-R strain). The brood stock was kept in 5 L glass tanks under flow-through conditions with dechlorinated tap water (hardness: 80 CaCO_3_ mg/L) at 25 °C under 16 h–8 h light–dark cycles. Fertilized eggs were collected from the stock, and the hatched larvae were maintained in the same conditions and fed freshly hatched, desalinated brine shrimp (*Artemia* spp.) nauplii until use.

Exposure to pyrene was performed according to the standard protocol of fish acute toxicity test (OECD 2019). Ten organisms (23-days-old) were transferred to a glass tank (19 cm × 4 cm × 20 cm height) containing 2 L of dechlorinated tap water with four replicates (i.e., a total of 40 organisms per treatment). In the exposure group, 200 μL of pyrene dissolved in acetone (1 g/L) was added to the water, resulting in a nominal concentration of 100 μg/L. The control group received 200 μL of acetone only. The concentration of 100 μg/L was selected as about one-tenth of reported 96 h LC50 value (1040 μg/L) (Ministry of the Environment 2009). The wet weight and total length of fish from the same cohort used in the toxicity test were 53.4 ± 8.7 mg and 15.2 ± 0.8 mm, respectively (*n* = 10, mean ± standard deviation), which meets the recommendation of the standard protocol (OECD 2019). During the exposure, the tanks were kept at 25 °C under 16 h–8 h light–dark cycles, and the fish were not provided with aeration or food. Whereas the measured pyrene concentrations were below the detection limit (the control) and 74 ± 2 μg/L (treatment, *n* = 4) at the start of exposure, they were below the detection limit (∼ 1 μg/L) in both groups at the end. The pyrene quantification was conducted by high-performance liquid chromatography following the method described in our previous study (Tani et al. 2021). The measured temperature, dissolved oxygen, pH, and conductivity were 24.8–25.4 °C, > 60% of saturation, 7.7–7.8, and 31 mS/m over 96 h, respectively. After 96 h of exposure, all fish in both the control and treatment groups were alive, so they were removed, euthanized on ice, and fixed with liquid nitrogen for transcriptome analysis of fish tissue. Just after removal of fish, 1.5 L of the water was filtered using a Sterivex filter cartridge with a nominal pore size of 0.45 μm (SVHV010RS, Merck) and a pump (Aqua Loader AL898, GL Science) at a filtration rate of 20–30 mL/min. The filter cartridge was then centrifuged at 3000 rpm for 5 min (S700T, Kubota) to remove water, and the resulting filter was closed with the caps (VRMP6 and VRSP6, ISIS Co., Ltd.) and subjected to eRNA extraction.

### 2.2. RNA extraction and sequencing

eRNA extraction was performed following a previous study (Tsuri et al. 2020) with slight modifications. Six hundred μL of RLT buffer containing β-mercaptoehanol was added to the filter cartridge and the filter was gently rotated for 1 hour at room temperature. The cartridge was placed on the 1.5 mL tube and centrifuged at 3000 rpm for 2 min (S700T, Kubota) to obtain the buffer. RNA in the buffer was extracted using RNeasy Mini Plus Kit (Qiagen) following the manufacturer’s protocol, but additionally performing DNase digestion on the RNeasy column membrane with RNase-free DNase set (Qiagen) for 30 min after wash by RW1 buffer. RNA was eluted in 50 μL water. For RNA from whole organisms, fish was homogenized in liquid nitrogen with a pestle. RNA was extracted from the homogenate using RNeasy Mini Plus Kit and treated with DNase in the same manner as eRNA, and pooled from two individuals for further sequencing.

RNA purity was verified at A260/A280 ratios larger than 1.9 using Nanodrop 1000 (Thermo Scientific). The RNA integrity number equivalent (RINe) was determined as 7.6 ± 0.4 (eRNA, range: 7.0–8.3) and 8.8 ± 0.6 (whole fish, range: 7.6–9.5) using TapeStation (Agilent Technologies). Although eRNA samples were somewhat degraded as indicated by lower RINe, they were used for subsequent analyses.

A cDNA library was constructed using TruSeq Stranded Total RNA Library Prep with Ribo-Zero Kit (Illumina) following the manufacturer’s protocol (1000000040499 v00), which eliminated ribosomal RNA (rRNA) and performed cDNA synthesis. Ribosomal elimination was performed instead of ploy-A transcript selection, because 3’ RNA degradation may occur in eRNA samples. The constructed library was sequenced on NovaSeq 6000 with a minimum 40 million read depth by Macrogen Japan (Tokyo). All raw reads were deposited on the Sequence Read Archive (accession number: DRR440008 –DRR440023 under the project PRJDB15208).

### 2.3. Data Analysis

The sequenced reads were mapped to the reference genome of *O. latipes* (accession number: GCF_002234675.1) using HISAT2 ver. 2.2.1 (Kim et al. 2019) with default settings. The mapped reads were counted using HTSeq ver. 0.11.3 (Anders et al. 2015) with the “-s reverse” option for strand-specific analysis, and the mapped reads were analyzed using R software ver. 4.2.1 and edgeR ver. 3.38.4 (Robinson et al. 2009; Chen et al. 2016). Genes with low read counts were removed using the filterByExpr function with the default settings to keep genes that have count-per-million (CPM) above 10 in 70% of fish tissue or eRNA samples (i.e, in at least 6 samples). Differentially expressed genes (DEGs) were determined using the estimateDisp, glmQLFit, glmQLFTest functions implemented in edgeR based on the criteria of a false discovery rate (FDR)-adjusted *p*-value < 0.1. Gene set enrichment analysis (GSEA) was performed using ShinyGO ver. 0.77 (Ge et al. 2020) based on the criteria of a FDR-adjusted *p*-value < 0.05. This analysis was conducted with all the detected genes for eRNA and fish RNA as background genes and included all available databases (e.g., Kyoto Encyclopedia of Genes and Genomes, Gene Ontology). Regression analysis of read counts was performed between eRNA and fish RNA, using a generalized linear model with a Poisson error distribution and a log link function. Principal component analysis (PCA) was performed using based on counts per million reads (CPM) using the prcomp R function with the “scale = TRUE” option. Heatmap for selected genes was created using ComplexHeatmap ver. 2.12.1 (Gu et al. 2016). The pattern of mapped reads on the reference genome was visualized using Integrative Genomics Viewer (IGV) ver. 2.8.2 (Thorvaldsdóttir et al. 2013).

Transcript integrity number (TIN) (Wang et al. 2016) was assessed as a metric for mRNA integrity using RSeQC ver. 2.6.4 (Wang et al. 2012). TIN measures mRNA integrity at transcript level based on mapped read coverage. To account for the bias due to RNA degradation especially for eRNA, the gene level read count was corrected using the TIN score and the loess R function, following the method by the previous study (Wang et al. 2016). When the corrected read counts became negative values, they were regarded as zero and then analyzed by edgeR.

To determine biological sources of unmapped reads, we performed taxonomic identification using DecontaMiner ver. 1.4 (Sangiovanni et al. 2019) with the database provided by the software’s web site (https://github.com/amarinderthind/decontaminer).

### 2.4. Quality assurance and quality control

Prior to the experiments, all equipment including glass tanks and pumps were treated with 10% bleach solution to remove DNA and thoroughly rinsed with Milli-Q water or exposure solution. The DNA contamination in eRNA and fish tissue RNA samples were checked by conventional PCR using primers PG17 for DMY gene (Matsuda et al. 2002) and 1% agarose gel electrophoresis.

## 3. RESULTS AND DISCUSSION

### 3.1. Overview of RNA-sequencing

For sixteen libraries, about 800 million reads were obtained with a length of 101 base pairs (see Tables 1 and S1). Of the sequenced reads, more than 92% of the fish tissue RNA were mapped to the reference genome, while 0.6 ± 0.4 % of the eRNA were mapped. This low mapping ratio for eRNA is comparable to 0.5% reported for eRNA of *D. pulex* (Hechler et al. 2022) and indicates that most of the sequenced eRNA originated from non-target microorganisms, such as bacteria (e.g., *Aeromonas, Shewanella*) and fungi (Figure S7).

**Table 1.** Summary of RNA-sequencing of whole fish and water samples.

|  | Fish |  | Water (eRNA) |  |
| --- | --- | --- | --- | --- |
|  | Control <sup>a</sup> | Pyrene | Control <sup>a</sup> | Pyrene |
| Number of sequenced reads (million) | 53.3 ± 7.5 | 44.2 ± 2.9 | 56.2 ± 8.0 | 45.2 ± 8.2 |
| Q30 (%) <sup>b</sup> | 95.5 ± 0.2 | 95.7 ± 0.2 | 95.7 ± 0.2 | 95.7 ± 0.1 |
| Ratio of mapped reads (%) | 93.7 ± 0.8 | 93.9 ± 0.7 | 0.8 ± 0.4 | 0.5 ± 0.2 |
| RNA integrity number equivalent (RINe) | 8.7 ± 0.8 | 8.8 ± 0.5 | 7.3 ± 0.3 | 8.0 ± 0.3 |
| Transcript integrity number (TIN) for mitochondrial genes | 91.6 (88.9–92.9) | 90.8 (89.0–92.5) | 89.1 (86.9–91.3) | 82.3 (76.1–88.0) |
| TIN for nuclear genes | 75.5 (60.5–83.4) | 75.6 (59.0–83.8) | 16.3 (7.8–29.7) | 10.2 (5.7–18.5) |

Mean ± standard deviation except median and inter-quartile range (25 and 75 percentiles) for TIN.
a: Solvent control containing 0.01% v/v acetone.
b: Ratio of bases that had phred quality score of over 30.

#### RNA degradation

RNA-seq analysis usually assumes that the amount of sequenced reads is proportional to the abundance and length of the transcript (Wang et al. 2016); however, this assumption can be distorted if RNA degrades. Despite all eight eRNA samples having RINe greater than 7.0 (Table S2 and Figure S1), the TIN values of eRNA were high enough for mitochondrial genes (median TIN: 83–91) but as low as < 20 of median for nuclear genes (Figure 1A). In contrast, both mitochondrial and nuclear genes in fish RNA had high TIN values (median TIN: > 73). These results were visually confirmed by genome browser, where genes with low TIN values showed skewed read coverage compared to genes with high TIN (Figure 1C, 1D). These results suggest that the commonly used metric of RNA integrity, RIN or RINe, may not accurately reflect the integrity of eRNA, which may contain rRNA originating from other species, as it relies on rRNA rather than mRNA.

**Figure 1.**
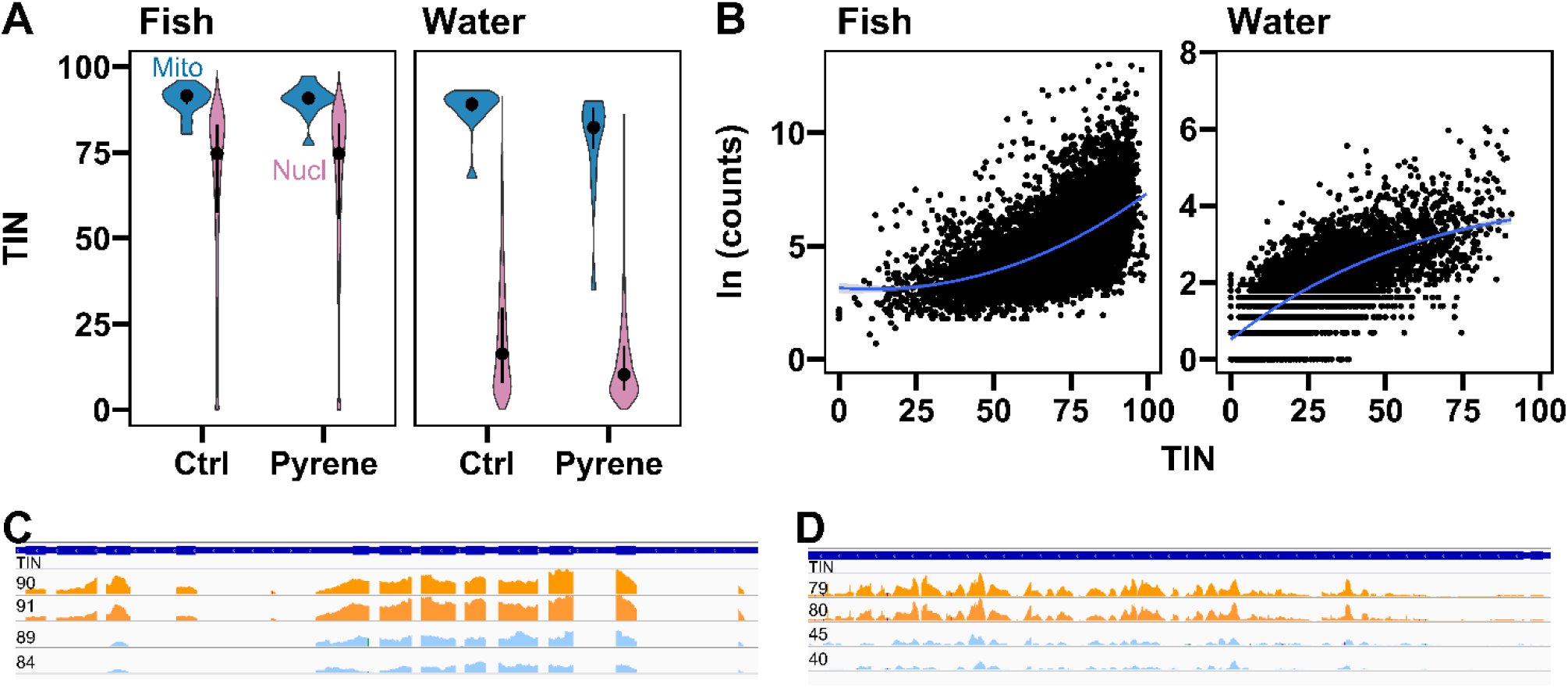
Association of transcript integrity number (TIN) with the source of RNA and read counts. Violin plots showing the distribution of TIN detected in fish RNA and eRNA (water), with different color representing mitochondrial (blue) and nuclear (pink) genes. (B) Examples of relationship between TIN and logarithmic raw read counts of fish RNA and eRNA in the control. Blue lines are locally estimated scatterplot smoothing (loess) lines. (C, D) Read mapping on the reference genome for *eff1a* (C) and *arnt* (D), as examples of genes with high and low TIN, respectively. Two fish (top, orange) and two water (bottom, blue) samples in the control are shown. TIN values are shown in left sides.

The lower TIN of nuclear eRNA observed in this study is consistent with previous findings of faster degradation rates for nuclear eRNA compared to mitochondrial eRNA (Marshall et al. 2021). Other studies have also reported faster degradation rates of nuclear eDNA compared to mitochondrial eDNA (Moushomi et al. 2019; Jo and Minamoto 2021), possibly due to the differences in protection afforded to DNA molecules (e.g, porousness of nuclear and mitochondrial membranes). However, a recent study showed comparable degradation rates between mitochondrial and nuclear eRNA (Jo et al. 2022).

The genes with low TIN values had low raw read counts in both fish RNA and eRNA (Fig 1B), indicating that RNA degradation could lead to biased read counts and affect the subsequent expression analysis. While a previous study have reported correlations between TIN and transcript length or GC content (Wang et al. 2016), we did not observe significant correlation in our data (Figure S2).

### 3.2. What genes were included in eRNA?

After filtering out genes with low expression, 1110 (eRNA) and 18,912 (fish tissue) genes were retained (Figure 2A), representing 4% and 70% of the *O. latipes* reference genome (26,845 genes), respectively. All genes detected in eRNA were also present in fish RNA. The genes in eRNA were involved in various biological functions, including ribonucleotide binding (GO:0032553), kinase activity (GO:0016301), and enzyme regulation (GO:0030234), and in several cellular components including ribosome (GO:0005840), cytosol (GO:0005829), cytoskeleton (GO:0005856), intercellular junction (GO:0005911). The median raw read counts for each gene were 197 (max: 452,462) and 15 (max: 4156) for fish tissue and eRNA, respectively.

**Figure 2.**
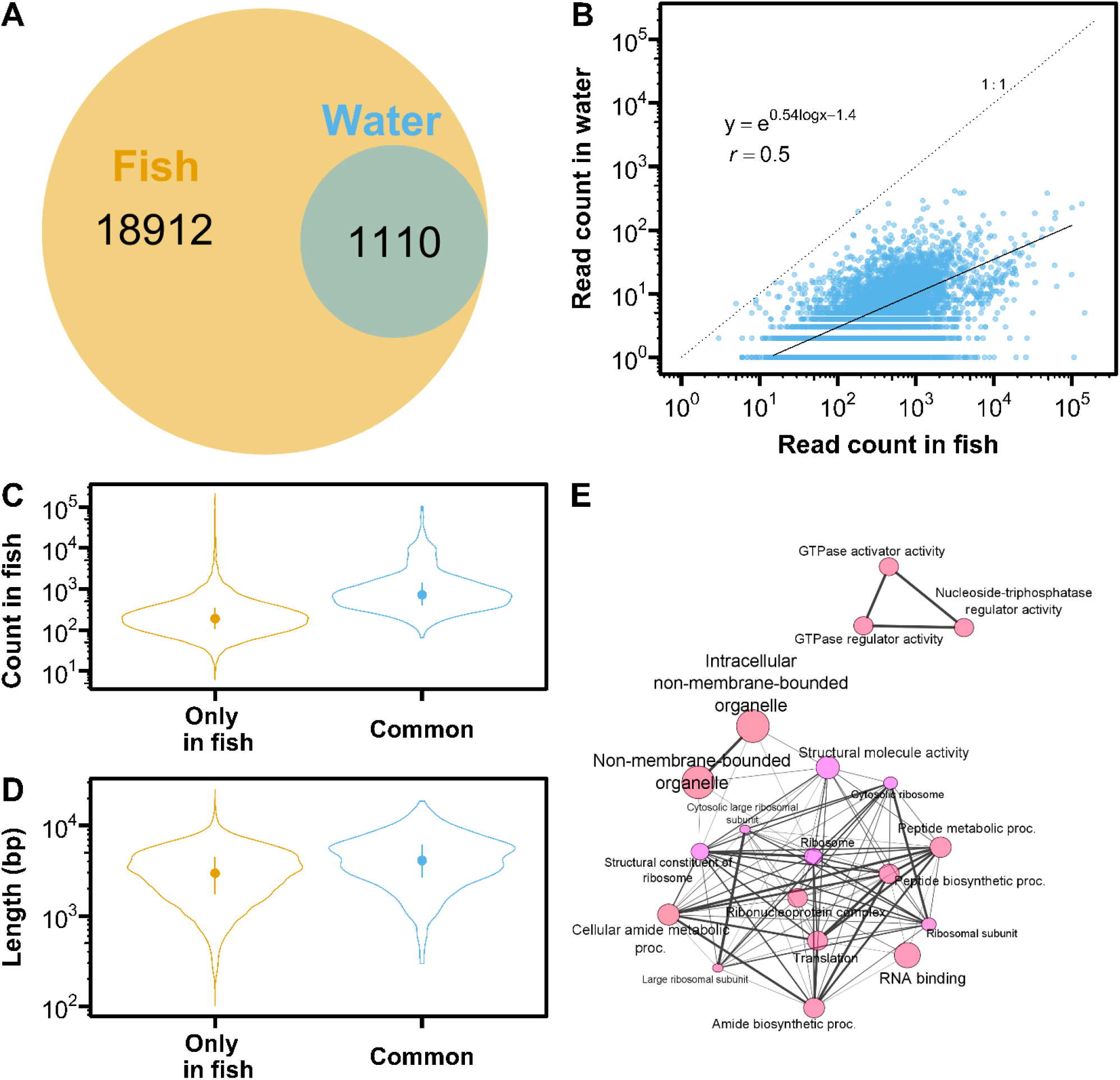
Comparison of expression profiles between fish tissue and environmental RNA (eRNA). (A) Venn diagram showing the number of genes detected in fish tissue and eRNA (water). (B) Correlation between raw read counts of fish tissue and eRNA. The dotted line indicates a 1:1 relationship, while the solid line is regression line with Poisson error distribution. Median read count was calculated for each gene from four control samples. (C) Comparison of raw read counts between eRNA and the genes specifically detected in fish tissue. Median read count for each gene was calculated from four control samples. Error bars indicate median and inter-quartile range (25 and 75 percentiles). (D) Comparison of transcript length between fish-specific and common genes. (E) Pathway networks enriched in eRNA compared to all genes detected in fish tissue. Vivid color and larger nodes are more significantly enriched (FDR-adjusted *p* value < 0.05) and larger gene sets, respectively. Edge widths represent the degree of gene overlap.

Expression levels of eRNA were correlated with those in fish tissue (Figure 2B, Pearson’s *r* = 0.50 on logarithmic scales). Additionally, in fish tissue, the genes shared between eRNA and fish tissue (“common” in Figure 2C) had higher expression levels (median counts: 698) than the genes found only in fish tissue (median: 183), suggesting that further deeper sequencing could detect a broader range of genes from eRNA. The common genes also had longer transcript length (median: 4102 bp) compared to the genes specific to fish tissue (median: 2962 bp) (Figure 2D), possibly due to the low sequencing-depth for eRNA, which can lead to biased sequencing toward long transcripts (Tarazona et al. 2011). There was no difference in GC content between the common and specific gene sets (median: 41.7% and 41.0%, respectively). Although GSEA showed that pathways related to ribosome (e.g., GO:0022626, GO:0003735) and GTPase activity (e.g., GO:0030695, GO:0005096) were significantly enriched in the gene sets shared between eRNA and fish tissue (Figure 2E), this enrichment might be due to the high expression levels in fish or long length of these common genes and not indicate that RNA involved in these pathways was selectively released from fish into surrounding water.

### 3.3. Can eRNA detect differential gene expression in fish?

As the results of differential expression analysis for pyrene-exposed fish and eRNA without TIN correction, we identified 311 genes (up: 196, down: 115) as differentially expressed in fish tissue and 86 genes (up: 46, down: 40) as differentially expressed in eRNA (Figure 3A). Among these DEGs, 6 up-regulated (LOC101165389, LOC101175077, LOC101165005, LOC101165259, *dsp*, and *fam83h*) and 6 down-regulated DEGs (*col1, col1a2*, LOC101173999, LOC101162163 [*col1a1b*], *socs3-2*, LOC101171531) were shared between fish tissue and eRNA, although enriched pathways were not shared (Tables S3 and S4). In eRNA, 74% of the DEGs showed the same directional change (up or down) as in fish tissue. Notably, three genes encoding collagen fibers were found to be shared between two RNA sources, and the related pathways (e.g., extracellular matrix structural constituent [GO:0005201], supramolecular fiber [GO:0099512]) were significantly enriched both in fish tissue and eRNA by pyrene exposure (Figures S3, S4). These results are consistent with a previous study showing a reduction in *col2a1* expression in rockfish embryos exposed to pyrene (Shi et al. 2012), and suggest that eRNA has the potential to detect changes in gene expression in fish tissue in response to chemical exposure.

**Figure 3.**
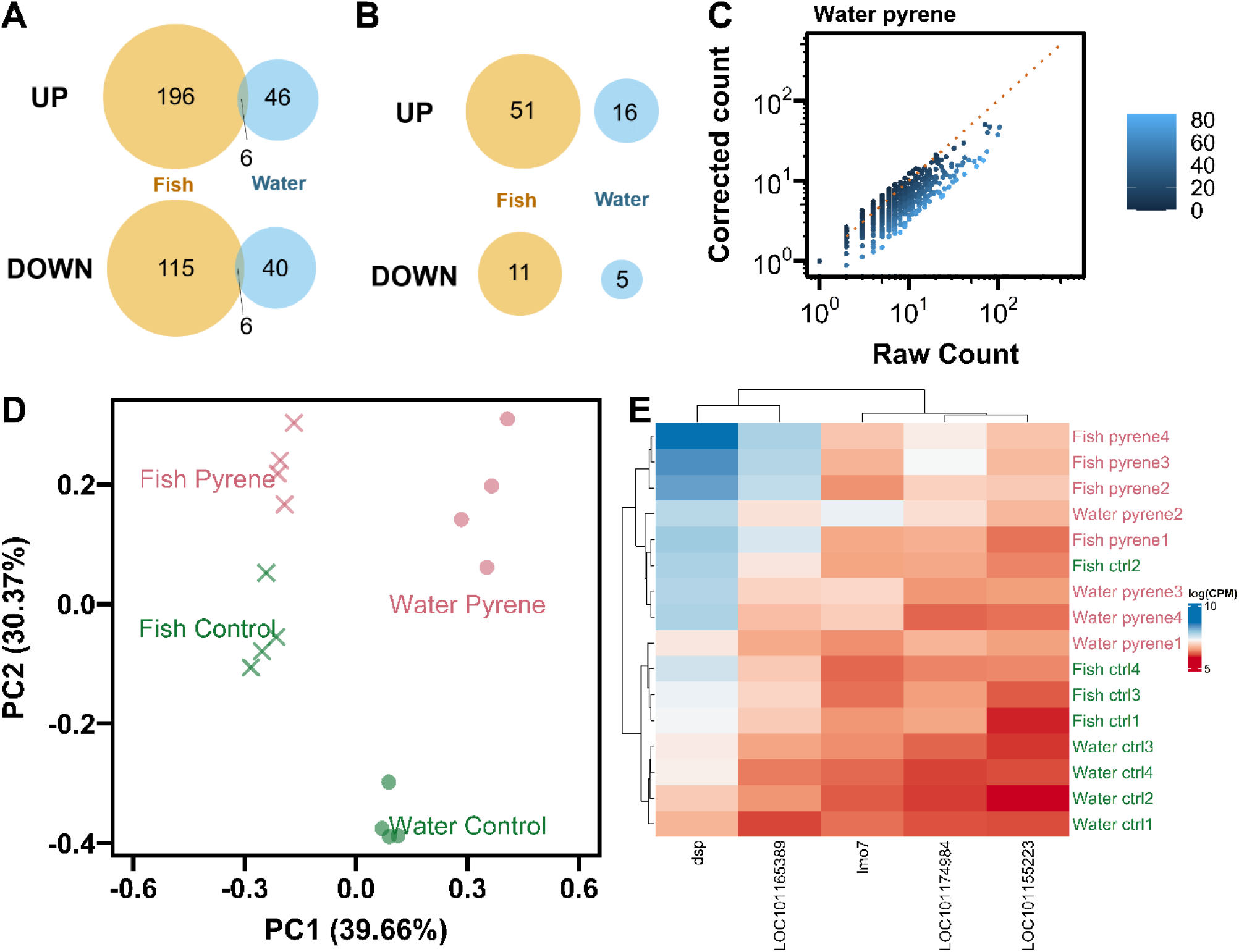
Differentially expressed genes (DEGs) in fish tissue and eRNA (water) in response to pyrene. Venn diagram showing the number of DEGs in fish tissue and eRNA, (A) without and (B) with TIN correction. (C) Example of comparison of read counts before and after TIN correction (for eRNA in the pyrene treatment). Color indicates TIN value of each gene. Dotted line is a 1:1 relationship. (D) Principal component analysis (PCA) based on the counts per million (CPM) of 86 DEGs in eRNA. PC1 and PC2 explained 39.66% and 30.37% of data variance. Different colors and symbols represent different exposure treatments and RNA sources, respectively. (E) Heatmap of 5 genes that had the largest loading scores for PC2. Color represents logarithmic CPM of each gene.

To determine if the low number of shared DEGs between fish tissue and eRNA could be attributed to RNA degradation in eRNA, we applied TIN correction. After TIN correction, read counts increased for highly degraded genes (i.e., low TIN) while those for lowly degraded genes decreased (Figure 3C). When using TIN-corrected read counts, the number of DEGs were reduced to 62 (up: 51, down: 11) for fish tissue and 21 (up: 16, down: 5) for eRNA (Figure 3B), and there were no DEGs shared between two types of RNAs. For fish tissue, DEGs and enriched pathways with TIN correction largely overlapped with those without TIN correction (Figures S3, S5, and S6). This indicates that TIN correction for fish tissue could provide biologically meaningful DEGs by accounting for RNA degradation and reducing the false positives, as demonstrated by a previous study (Wang et al. 2016). In contrast, for eRNA, DEGs with TIN correction did not overlap with those without TIN correction (Figure S6), possibly due to the low sequencing-depth in eRNA being too low to accurately reflect the actual gene expression levels after TIN correction. In fact, negative (which was transformed to zero before edgeR analysis) and quite small read counts were observed in eRNA samples after TIN correction (Figure 3C). Therefore, TIN correction was not performed in further analyses in this study.

Although we could not identify the cause of the low number of shared DEGs, we suggest that this should be re-examined using sufficient sequencing depth.

PCA based on the CPM of 86 DEGs in eRNA showed that the samples were clearly separated by RNA sources along the first principal component (PC1) accounting for about 40% of overall variance (Figure 3D). Additionally, although not completely, the samples were separated by pyrene exposure along the second principal component (PC2), which was attributed to genes with the largest loading scores for PC2, including *dsp, lmo7*, LOC101155223, LOC101174984, and LOC101165389 (Figure 3E). Notably, two of these genes (*dsp* and LOC101165389) were commonly identified as DEGs between fish tissue and eRNA. These results suggest that eRNA can be a useful non-invasive tool for detecting molecular changes in response to chemical exposure in fish.

### 3.3. Limitation and recommendation

The use of non-invasive tools for assessing physiological status in organisms is becoming increasingly important due to growing concerns about animal welfare. eRNA represents a promising avenue for researchers in research fields of ecology and ecotoxicology in this regard. As demonstrated by this study and recent papers (Tsuri et al. 2020; Hechler et al. 2022), eRNA contains genes involved in a wide range of biological processes and molecular functions and it was able to detect differential gene expressions in response to environmental stressors. While these findings are novel in ecology and ecotoxicology, they are not surprising given that a variety of RNAs have been detected in human sweat and urine (Merchant et al. 2017; Bart et al. 2021). The detection of general stress- and defense-related genes, including catalase (*cat*), superoxide dismutase 1 (*sod1*), *sod2*, glutathione peroxidase 7 (*gpx7*), heat shock protein 90 (*hsp90ab1*), and aryl hydrocarbon receptor nuclear translocator (*arnt*) (Figure 4), further supports the potential of eRNA as a non-invasive tool for assessing the health status of aquatic organisms. Moreover, the experimental design in this study (e.g., fish density, water volume) aligned with the standardized toxicity test protocol (OECD 2019), highlighting that eRNA analysis does not require an unrealistic large volume of water or high fish density and can be applicable to routine toxicity tests. This suggests that the combination of eRNA analysis and flow-through exposure system could enable time-course gene expression profiling in a non-invasive manner.

**Figure 4.**
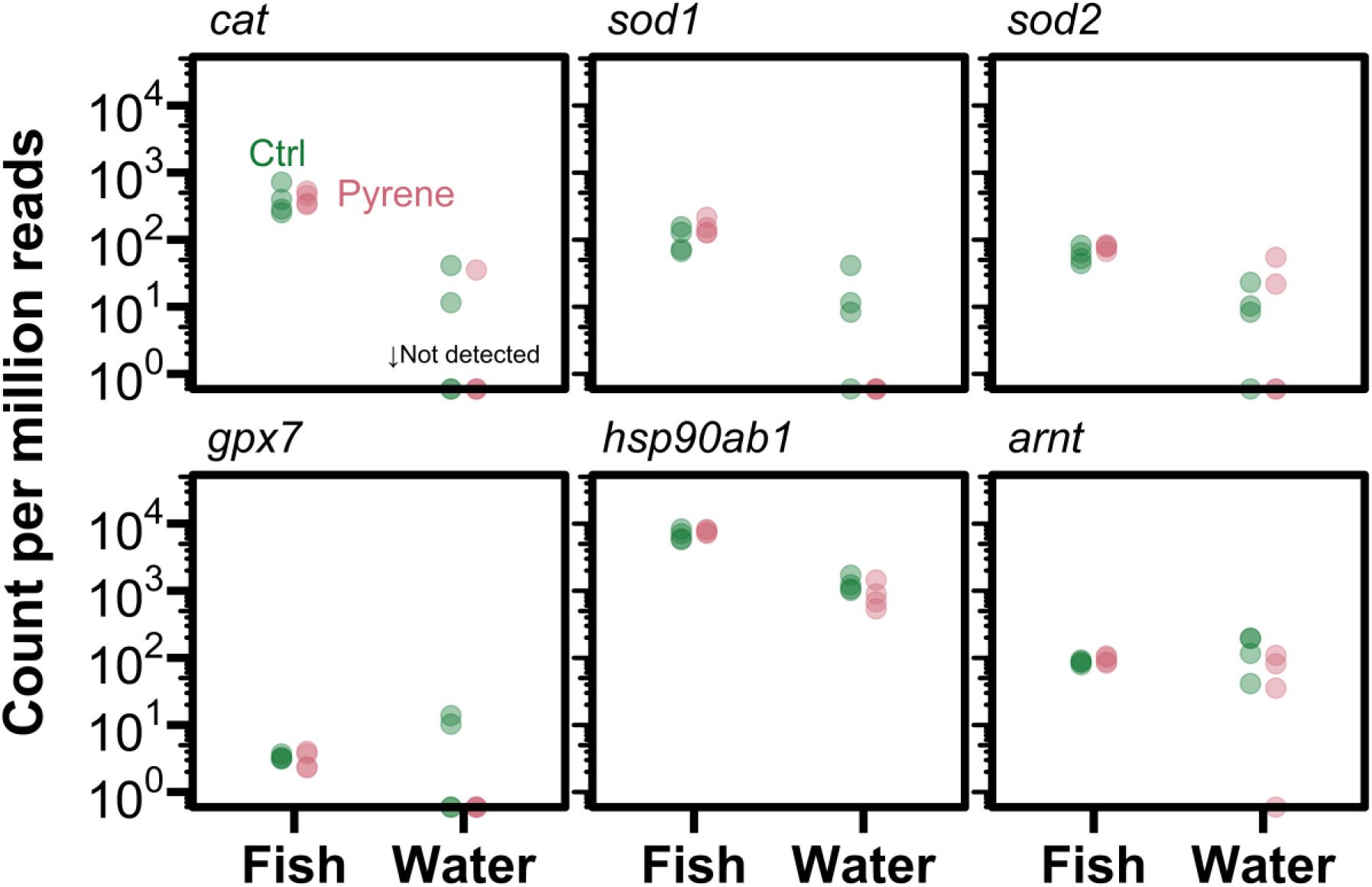
Counts per million reads of selected stress- and defense-related genes in fish RNA and eRNA. Dots on the x axis indicate below the detection limit. Different colors represent different exposure conditions. TIN correction was not performed.

Despite the potential of eRNA, this study also highlighted some of its limitations. Firstly, eRNA extracted from surrounding water contained RNA from non-target (micro) organisms, which resulted in quite low sequencing-depth and biased sequencing of longer transcripts, leading to a low number of DEGs identified. Although few million mapped reads ware sufficient to identify and quantify highly expressed genes, > 40–80 million reads are required to quantify lowly expressed genes and to perform differential expression analyses (Sims et al. 2014). In this study, the sequencing depth in eRNA fell short of this standard by < 1%. To overcome this issue, we recommend that future study should perform targeted RNA-Seq (Wang et al. 2018) or quantitative PCR to increase the reliability of eRNA quantification. Such improvements may reveal the difference in expression profiles between eRNA and organism tissue RNA, which could not be clarified in this study due to the low sequencing-depth.

For example, the expressions of genes involved in skin and mucus may be dominant in eRNA. The sequencing improvements may also help the application of eRNA analysis to natural environments, where coexisting RNA from non-target organisms is greater than in laboratory conditions. Secondly, we showed that eRNA, particularly nuclear gene, was highly degraded. The degree of degradation varied between genes, indicating the need for correction of expression levels. Although we applied the TIN correction method to eRNA, the correction might not be successful due to the low sequencing-depth in eRNA. However, we hope that the correction method applied in this study will be helpful in future eRNA studies with a large sequencing-depth. Additionally, we recommend exploring other correction methods proposed in recent studies (Xiong et al. 2019) to improve the accuracy of eRNA applications.

## ACKNOWLEDGMENTS

The authors are grateful to H. Takahashi for her help in culturing test organisms. This study was financially supported by National Institute for Environmental Studies (NIES) Research Funding (Type B).

