## Supplementary file for "Environmental RNA as a non-invasive tool for assessing toxic effects in fish: a proof-of-concept study using Japanese medaka exposed to pyrene"

### List of materials

Table S1–S5
Figure S1–S7
References

Table S1. Summary of RNA-sequencing.

| | | Number of sequenced reads (million) | Q30 (%) | Ratio of mapped reads (%) |
|---|---|---|---|---|
| Fish | Control 1 | 55.8 | 95.5 | 94.12 |
| | Control 2 | 42.2 | 95.5 | 94.27 |
| | Control 3 | 56.2 | 95.6 | 92.60 |
| | Control 4 | 58.8 | 95.2 | 93.66 |
| | Pyrene 1 | 40.9 | 95.5 | 93.01 |
| | Pyrene 2 | 42.8 | 95.7 | 94.05 |
| | Pyrene 3 | 47.4 | 95.9 | 94.69 |
| | Pyrene 4 | 45.8 | 95.6 | 93.98 |
| Water | Control 1 | 61.1 | 96.0 | 0.68 |
| | Control 2 | 55.3 | 95.5 | 0.54 |
| | Control 3 | 63.0 | 95.5 | 0.58 |
| | Control 4 | 45.2 | 95.6 | 1.42 |
| | Pyrene 1 | 52.2 | 95.7 | 0.22 |
| | Pyrene 2 | 45.5 | 95.7 | 0.56 |
| | Pyrene 3 | 43.4 | 95.6 | 0.77 |
| | Pyrene 4 | 50.9 | 95.7 | 0.34 |

Table S2. Details of RNA degradation indices.

| | | RNA integrity number equivalent (RINe) | Median transcript integrity number (TIN) | |
|---|---|---|---|---|
| | | | Mitochondria | Nuclear |
| Fish | Control 1 | 7.6 | 91 | 77 |
| | Control 2 | 9.4 | 91 | 77 |
| | Control 3 | 8.9 | 89 | 75 |
| | Control 4 | 8.9 | 93 | 79 |
| | Pyrene 1 | 8.8 | 89 | 73 |
| | Pyrene 2 | 9.5 | 91 | 78 |
| | Pyrene 3 | 8.7 | 90 | 77 |
| | Pyrene 4 | 8.3 | 91 | 79 |
| Water | Control 1 | 7.4 | 90 | 28 |
| | Control 2 | 7.2 | 90 | 24 |
| | Control 3 | 7.6 | 90 | 26 |
| | Control 4 | 7.0 | 91 | 30 |
| | Pyrene 1 | 7.7 | 84 | 19 |
| | Pyrene 2 | 8.1 | 89 | 23 |
| | Pyrene 3 | 8.3 | 88 | 27 |
| | Pyrene 4 | 7.7 | 83 | 18 |

Table S3. Top six pathways in fish RNA enriched by pyrene exposure, without transcript integrity number (TIN) correction.

| Enrichment FDR | No. of genes | No. of total genes in pathway | Fold enrichment | Pathway |
|---|---|---|---|---|
| 0.001 | 7 | 73 | 13.8 | Extracellular matrix structural constituent |
| 0.005 | 3 | 3 | 60.9 | DNA photolyase activity |
| 0.025 | 22 | 862 | 2.6 | Response to external stimulus |
| 0.025 | 3 | 5 | 36.5 | Cone photoresponse recovery |
| 0.029 | 8 | 87 | 5.8 | ECM-receptor interaction |
| 0.034 | 5 | 42 | 10.5 | Phototransduction |

Enrichment analysis was performed using ShinyGO (ver. 0.77) (Ge et al. 2020).

Table S4. Top two pathways in environmental RNA (eRNA) enriched by pyrene exposure, without transcript integrity number (TIN) correction.

| Enrichment FDR | No. of genes | No. of total genes in pathway | Fold enrichment | Pathway |
|---|---|---|---|---|
| 0.004 | 5 | 28 | 9.9 | Intermediate filament-based process |
| 0.004 | 5 | 28 | 9.9 | Intermediate filament cytoskeleton organization |
| 0.004 | 7 | 76 | 6.5 | Intermediate filament |
| 0.004 | 7 | 77 | 6.5 | Intermediate filament cytoskeleton |
| 0.004 | 9 | 431 | 4.8 | Supramolecular polymer |
| 0.004 | 9 | 430 | 4.8 | Supramolecular fiber |
| 0.026 | 9 | 528 | 3.8 | Supramolecular complex |
| 0.028 | 3 | 5 | 13.9 | Intermediate filament binding |

Enrichment analysis was performed using ShinyGO (ver. 0.77) (Ge et al. 2020).

Table S5. Top eight pathways in fish RNA enriched by pyrene exposure, considering transcript integrity number (TIN) correction.

| Enrichment FDR | No. of genes | No. of total genes in pathway | Fold enrichment | Pathway |
|---|---|---|---|---|
| 0.002 | 4 | 73 | 39.3 | Extracellular matrix structural constituent |
| 0.002 | 5 | 87 | 18.1 | ECM-receptor interaction |
| 0.017 | 6 | 247 | 7.7 | Focal adhesion |
| 0.017 | 5 | 265 | 9.9 | External encapsulating structure |
| 0.017 | 5 | 265 | 9.9 | Extracellular matrix |
| 0.028 | 2 | 9 | 76.1 | Desmosome |

Enrichment analysis was performed using ShinyGO (ver. 0.77) (Ge et al. 2020).

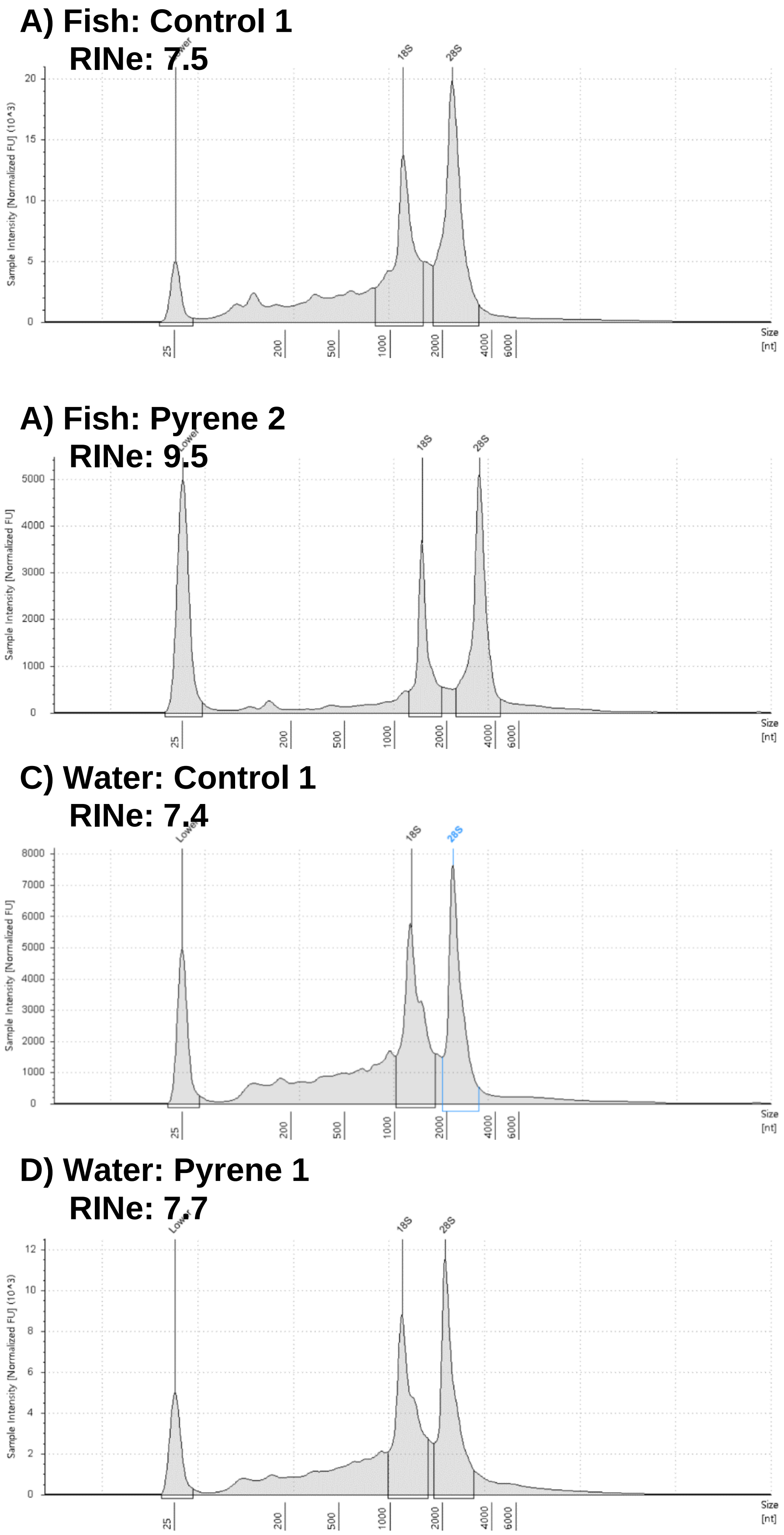


Figure S1. Examples of total RNA electropherogram of fish RNA and eRNA by TapeStation. A) Fish RNA in the control. B) Fish RNA in the in the exposure treatment. C) Water eRNA in the control. D) Water eRNA in the exposure treatment.

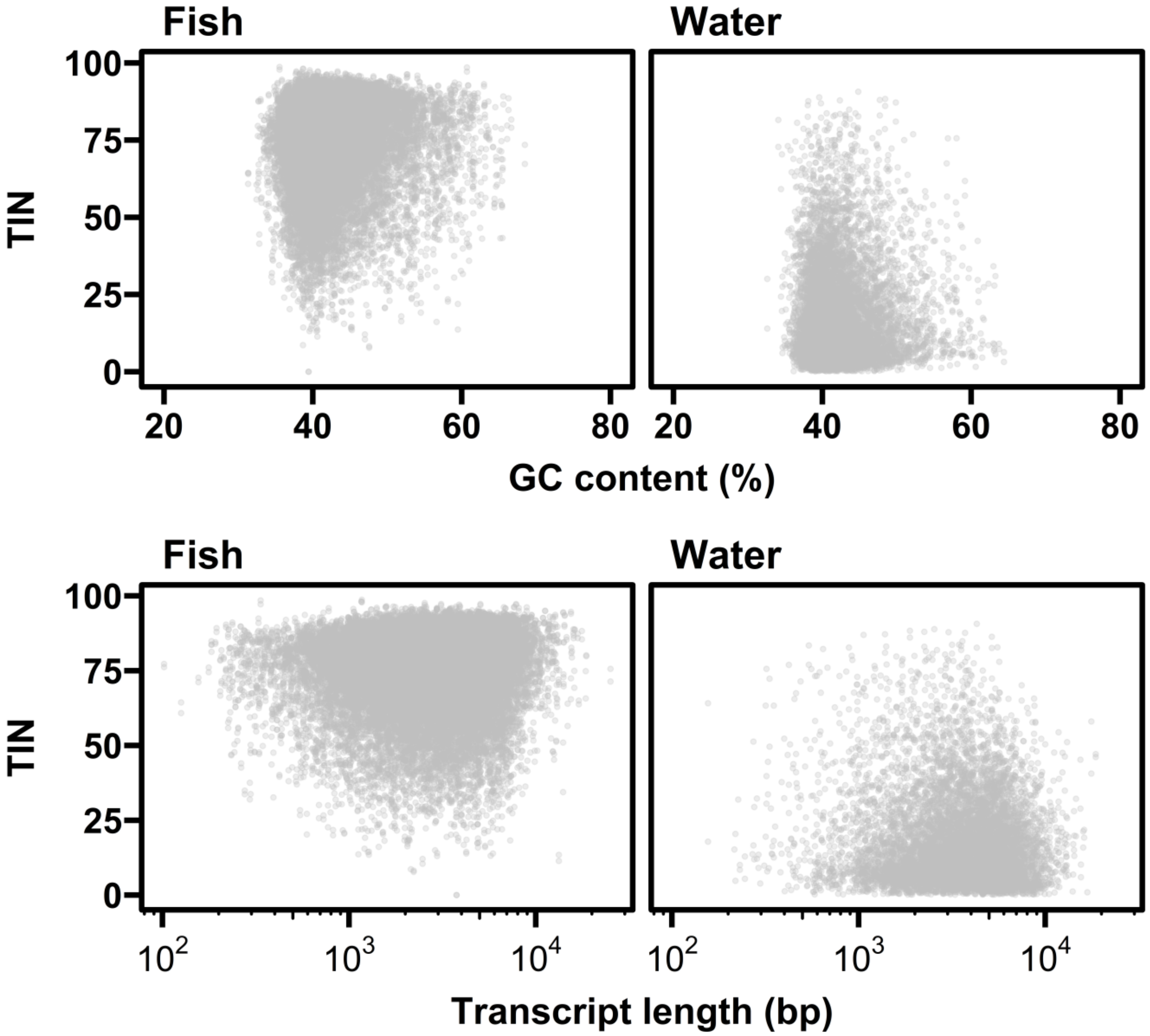


Figure S2. Relationship between transcript integrity number (TIN) and transcript length or GC content.

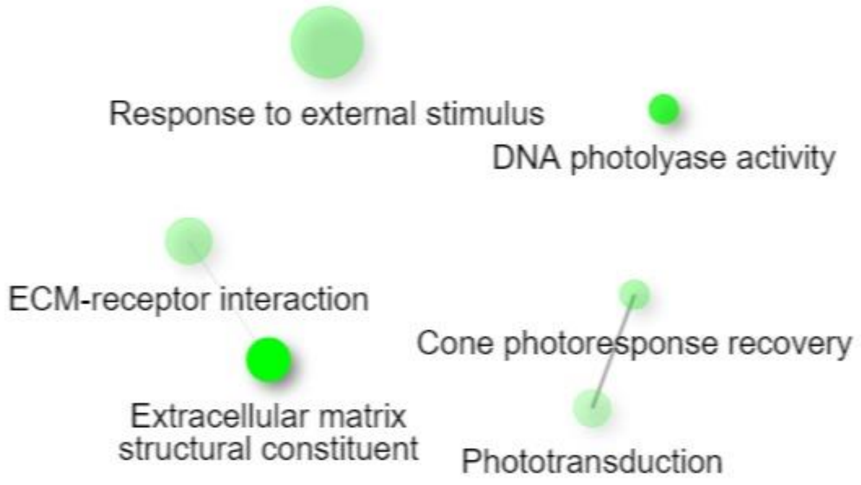


Figure S3. Pathway networks in fish RNA enriched by pyrene exposure, without transcript integrity number (TIN) correction. Darker and larger nodes are more significantly enriched (FDR-adjusted p value < 0.05) and larger gene sets, respectively. Edges represent the degree of gene overlap. Enrichment analysis was performed using ShinyGO (ver. 0.77) (Ge et al. 2020).

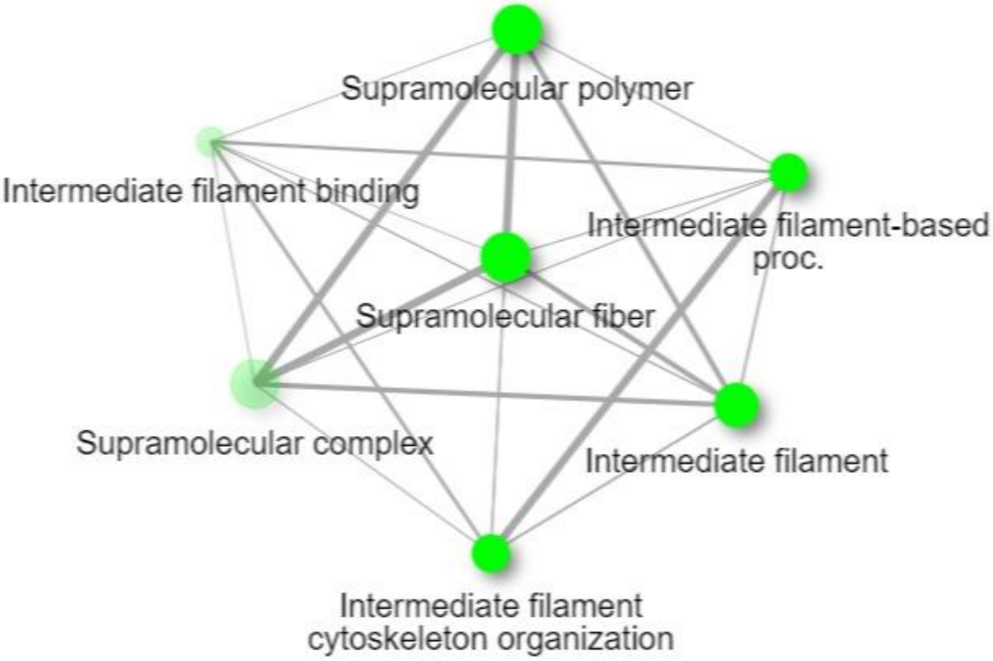


Figure S4. Pathway networks in eRNA enriched by pyrene exposure, without transcript integrity number (TIN) correction. Darker and larger nodes are more significantly enriched (FDR-adjusted p value < 0.05) and larger gene sets, respectively. Edges represent the degree of gene overlap. Enrichment analysis was performed using ShinyGO (ver. 0.77) (Ge et al. 2020).

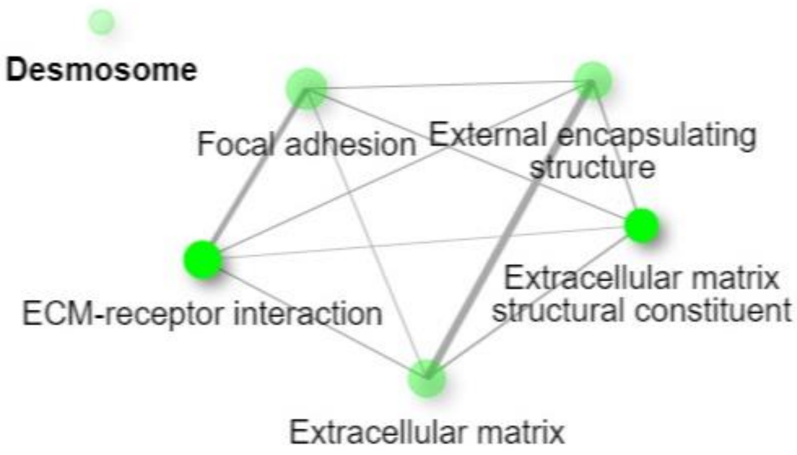


Figure S5. Pathway networks in fish RNA enriched by pyrene exposure, considering transcript integrity number (TIN) correction. Darker and larger nodes are more significantly enriched (FDR-adjusted p value < 0.05) and larger gene sets, respectively. Edges represent the degree of gene overlap. Enrichment analysis was performed using ShinyGO (ver. 0.77) (Ge et al. 2020).

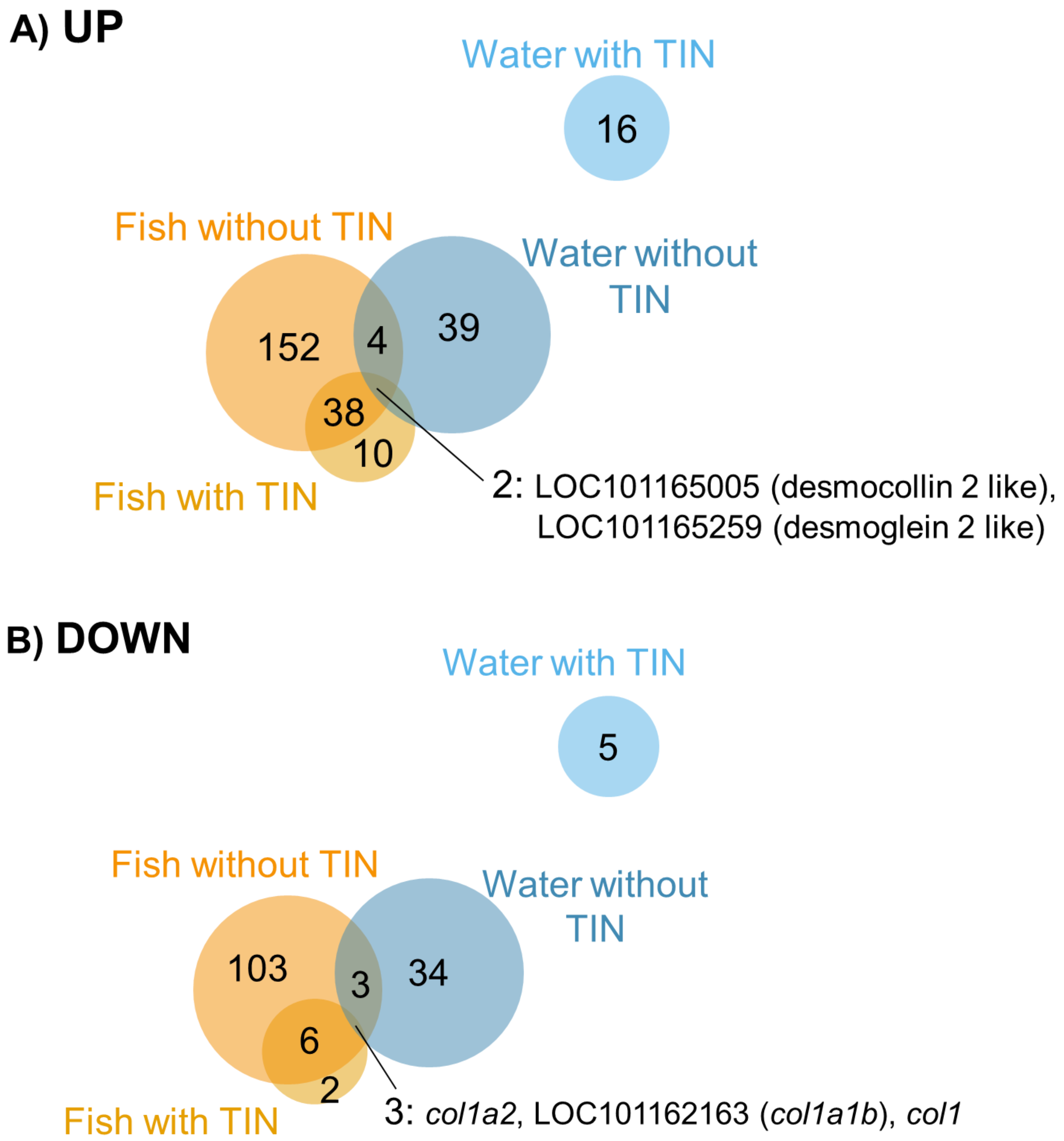


Figure S6. Venn diagram showing the overlap of differentially expressed genes (DEGs) by pyrene exposure with and without transcript integrity number (TIN) correction.

**(A) Bacteria**

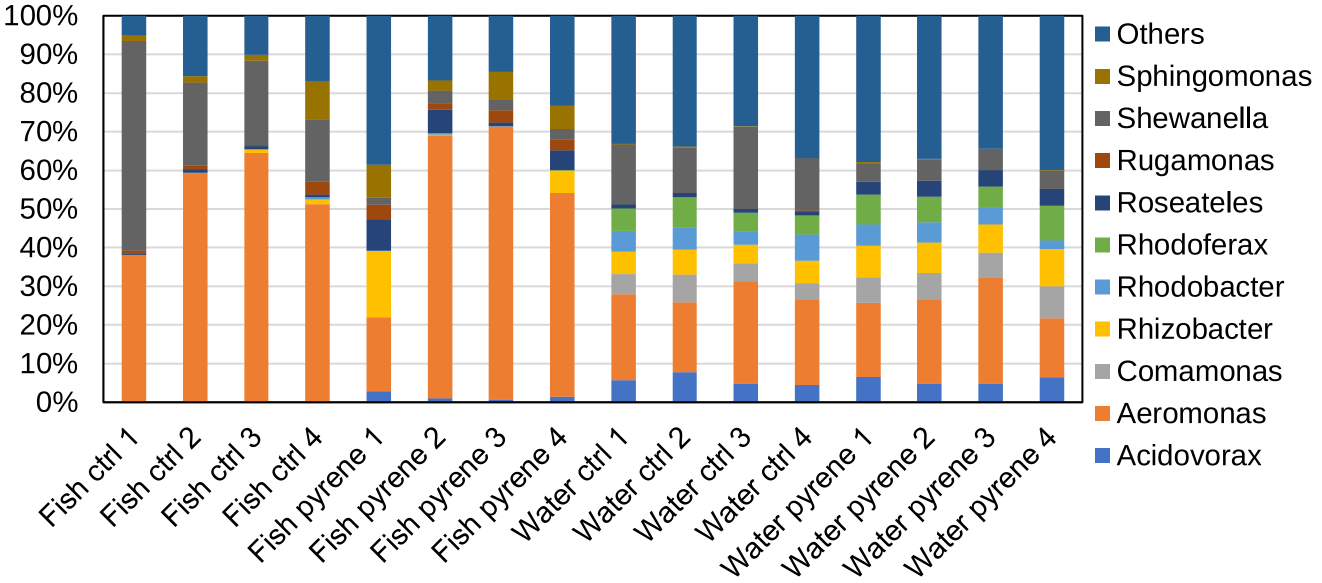


**(B) Fungi**

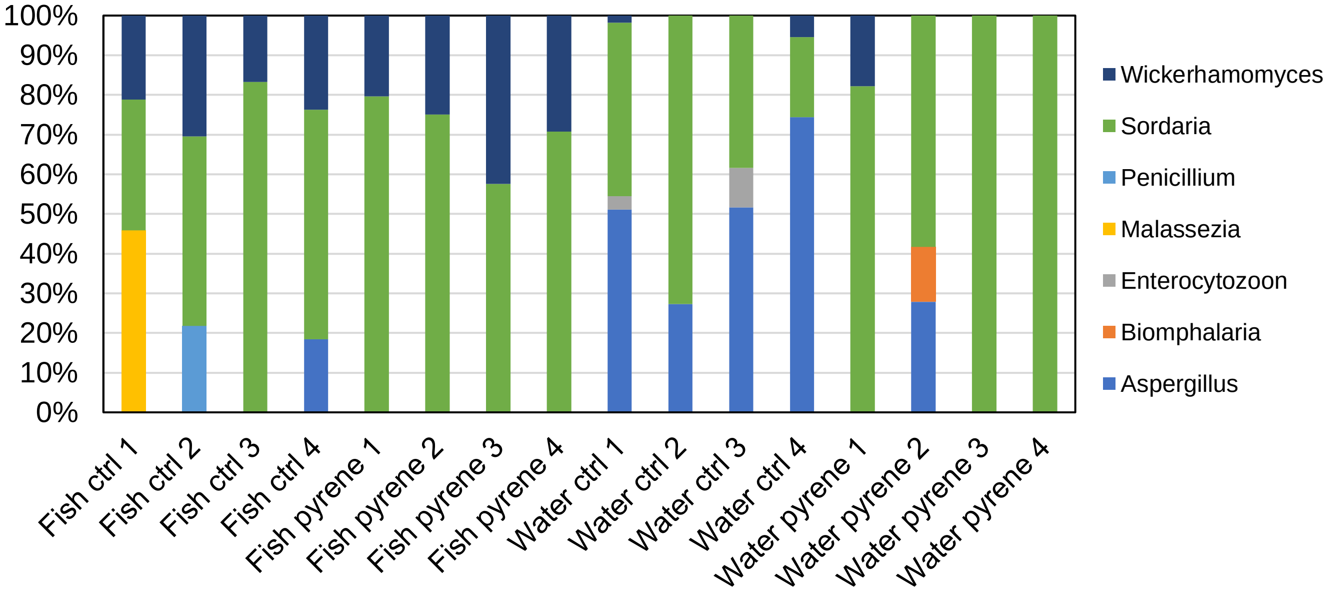


Figure S7. Relative abundance of (A) bacteria and (B) fungi at genus levels in fish tissue and water RNA samples. Taxonomic identification was performed using DecontaMiner (Sangiovanni et al. 2019).
